# The free energy of an ecosystem: towards a measure of its inner value

**DOI:** 10.1101/2022.12.19.521149

**Authors:** Gerardo M. E. Perillo, Mariana I. Zilio, Fernando Tohme, M. Cintia Piccolo

## Abstract

Based on a free energy approach, we propose the estimation of ecosystem Inner Value, which is both non-instrumental and objective, reflecting the ecosystem’s value for itself as a natural entity, abstracted from any human valuation activity. The ecosystem services approach has become the dominant criteria for studying the relationship between humans and nature. Although there is concern about preserving and recuperating damaged ecosystems, we seldom consider how much the ecosystem values itself. Then, we propose that Inner Value could be a tool to evaluate and model ecosystems’ health before any anthropic disturbance, allowing comparison with the impact these disturbances may have in the future. We also suggest that it should be a requirement for any Environmental Impact Assessment.

**Significance Statement:** Ecosystem are valued for the services they provide to human. Therefore, they are estimated following human (mostly economic) criteria. However, any ecosystem has its own value independent of the services it can give. Therefore, we propose the use of the Inner Value concept as a non-instrumental, objective measure reflecting the ecosystem’s value for itself as a natural entity based on a free-energy approach. Inner Value could be a tool to evaluate and model ecosystems’ health before any anthropic disturbance, allowing comparison with the impact these disturbances may have in the future.

## Introduction

The value of nature has been (and still is) the object of a profound debate that involves philosophical, moral, and ethical discussions (1). Each value category is derived from a different cognitive framework that reflects a given relationship with nature (2). There is a consensus that the biosphere has value *per se* but, what kind of value? related to what? or even for whom? (3) are still questions extensively debated.

One of the most significant and widespread research field aimed at studying the relationship between humans and nature is the Ecosystem Services Science (4) mainly focuses on defining, classifying (5–7), and estimating the value of the services they provide (8—10) as well as to define and assess the risks they are facing (11–13). Risks are generally due to changing global, regional, and local conditions and anthropic activities as well, while valuation depends primevally on the notion of value adopted.

Of course, there is plenty of concern about preservation and even recuperation of damaged or fragmented ecosystems, but with a psychological sensation (e.g., culpability?) that humans are a species that tends to destroy the planet for its benefit, regardless of the consequences. Seldom, if ever, we think from the ecosystem point of view. What the ecosystem needs or wants for itself or, more importantly, how much value the ecosystem “thinks” it has itself.

In this context, we propose a new paradigm by defining a mechanism of valorization of the ecosystem using a completely different approach and unit of measure than those used by the specific literature hitherto. Therefore, we define the Inner Value of an ecological network as a normalized amount of the free energy concept (14). Our objective is to find a measure of the value of an ecosystem, both non-instrumental and objective, which reflects the value that the ecosystem has for itself as a natural entity, abstracted from any human valuation activity.

### Value and valuation of nature

The instrumental value is perhaps the more explored and discussed value of ecosystems: nature has value because it plays a role in supporting and furthering human interests or, in other words, the value of nature consists in what it can do for us (1). Instrumental covers a variety of values: use-value (consumptive and non-consumptive, direct or indirect); option values (option value proper and quasi-option value); and non-use values. Among the latter, which reflects the willingness to pay to preserve the environment for the benefit of other people, intra and intergenerationally (15), the more studied is the existence value motivated by altruism, heritage, symbolism, or the belief in the right to the existence of other forms of life: a position congruent with the different variants of non-anthropocentric ethics (3).

Except for the value related to the last two reasons, all instrumental values can be considered extrinsic values because what is valued is something other than the specific good, for our own well-being or for the well-being of others. Except for the value related to the last two reasons, all instrumental values can be considered extrinsic values, because what is valued is something other than the specific good, for our own well-being or for the well-being of others.

On the contrary, intrinsic values refer to an essential type of value, given by the contribution of an object or action to maintain the health and integrity of an ecosystem or species per se, no matter any human satisfaction (16). Then, intrinsic value tries to capture the value attached to the environment and life forms for their own sake (15), beyond their ability to fulfil human needs.

Despite this apparent simplicity, the notion of intrinsic value is still confused and widely debated in the literature (17–20). The discussion about instrumental and intrinsic value have also generated new developments, mainly aimed to improve the environmental policy. Thus, the relational value concept (17) emerges as a third kind of value, which is not present in things but reflects the relationships and responsibilities to them. For instance, Rowlands (1) identified three different interpretations of intrinsic value: the value that an object possesses in virtue of its intrinsic properties or features; the value understood as non-instrumental value; and the value that objects have irrespective of whether sentient creatures value them or not. Davidson (21) defines warm glow value and existence value as the satisfaction that people derive from altruism towards nature and the mere knowledge that nature exists, respectively, and differentiates such values from intrinsic value, reflecting the value that nature has for itself.

This definition implies that intrinsic value lies outside of the ecosystem services framework. Nevertheless, such exclusion does not necessarily imply that ecosystem services are an anthropocentric approach in the moral sense since it does not deny that nature has value for itself. Instead, the intrinsic value lies outside because it does not reflect nature’s value from a human perspective (or, in Rowlands’ (1) terms, its instrumental use for all sentient creatures).

Instrumental and intrinsic values reflect different aspects of what nature does for itself or us, respectively. Nevertheless, in both cases, the perspective from which an environmental good or service is valued is purely and exclusively anthropocentric, in the sense that it is impossible to dissociate the value from the activity of valuation made by individuals.

The ecosystem services framework, undoubtedly the dominant paradigm for describing the relationship between humans and nature in our times, considers that the ecosystem functions become services only if they satisfy people’s needs and increase their welfare. Then, the ecosystem services provision depends on the degree of dependence of those services by certain groups and the relationship of such a service with other ecosystem services. In the equation, there is little or no focus on the conditions or potentialities of the ecosystem other than that required to maintain the ecosystem as a service provider for satisfying human needs, which means that instrumental value prevails all over other types of values. Then nature has no value besides human valuation.

Logically, valuation is always a subjective basis. Consequently, there is not a unique value for each ecosystem service, which generates a wide range of interpretations when judging the potential of, for instance, preservation of the ecosystem or extraction of products (either tangible or intangible). The valuation also may vary as a function of time (22, 23). The rate of variation of the ecosystem value depends on the relative scarcity of the service, the need that the service satisfies, and the use it will have rather than the actual value that the service has per se. Also, the ecosystem’s valuation is based on the services it can provide depending on whether it has some particular value in the past, present, or future. These values clearly vary with time, even if they are essential, considering the market estimation of the service, and even can vary for different cultural or regional approaches.

The lack of consensus about how an ecosystem (whatever its nature) must be assessed has created a deep void among most economists. They are trying to evaluate the services in monetary terms, and ecologists concerned about the system’s past, present, and future status. To abstract from such extreme positions and avoid the usual anthropogenic logic underlying the valuation of nature, we propose the notion of Inner Value derived from both the idea of ecological networks and the need to find a non-instrumental and objective measure of the value of a system.

### Ecological Networks

Ecosystems, even those that we may consider as simple ones, are characterized by a complex spatial heterogeneity of their features and their services (5). To develop their complete life cycle, individual elements of an ecosystem require several interactions with the other (but not necessarily all) members of the ecosystem. The energy transfer can describe those interactions from one individual taxon to another (24, 25) and their relationship with the surrounding structure (i.e., geomorphology, nutrient input/output) and external input/output (i.e., solar radiation, gas exchange).

Although we do recognize the importance of the exchange with the exterior, for the sake of our analysis, we consider the ecosystem as a closed system receiving only solar radiation and nutrient input from the sediment surrounding the system. Our idea is to track and evaluate how the interaction of the taxa based on the ecological trophic transfers provides an estimation of the Inner Value of the whole ecosystem at a specific time, as well as how we can use the Inner Value as a tool to analyze the ecosystem self-evolution over time. An ecosystem is integrated by several biological organisms aggregated in taxa and the surrounding physical (abiotic) conditions as a physical environmental structure. Organisms interact among them, and the various factors that continually influence their behavior and evolution.

Another way to see an ecosystem is like an ecological network where the interaction among taxa and the surrounding environment is based on a transfer of energy, and this transfer can be either in one direction or bidirectional. The fact that the same external factors may interact with different taxa at different rates and the various interaction mechanisms that the biotic organisms have among themselves analyzes ecological networks as rather complex but also variable at different times of the ecosystem life. In other words, the same exchange between two particular taxa may be different depending on the degree of maturity of the ecosystem and may change if the mass of one taxon has different values at different stages of, for instance, a seasonal cycle. Furthermore, some taxa may interact with another under some circumstances, but if the appropriate conditions are not met, the interaction may have a different magnitude or even be null.

Since we assume the ecosystem is closed, we are not considering any influence of climatic variables and water input from rivers or groundwater. We intend to evaluate the energy exchange among the various ecosystem components towards obtaining its inner value. We then define the Inner Value *as a measure of how the elements of the ecosystem determine the significance of the ecosystem by itself before any exchange with the external world*.

Basically, we look to separate the ecosystem from its external factors to maximize the inner value. Then, once the system is open to the exterior, we could evaluate how the input/output may affect this inner value and its evolution. Furthermore, the inner value could also be employed as a measure of the internal efficiency of the system’s organization and, by continuously maximizing it, developing tools for mitigation or resilience.

### Inner Value from Free Energy

Complex systems can either fail or succeed based on how they resist internal and external tendencies towards their disintegration. This is the case of ecological networks, which are as self-organizing systems. That is, systems in which their internal dynamics contribute to preserve their structure and behavior. Lack of thermodynamical equilibrium with its environment provides a clear indication of live activity.

All ecosystem processes are irreversible (26). At each process, energy is lost in the form of heat that contributes to entropy production (27), implying that processes involving energy flux are associated with the disorder (28). There are two primary types of incoming energy (Fig. 1): (i) solar energy that enters the ecosystem via photosynthesis and (ii) the energy bound in chemical components which flows through the boundaries of ecosystems via inorganic or organic nutrients pools (29).

**Figure 1.**
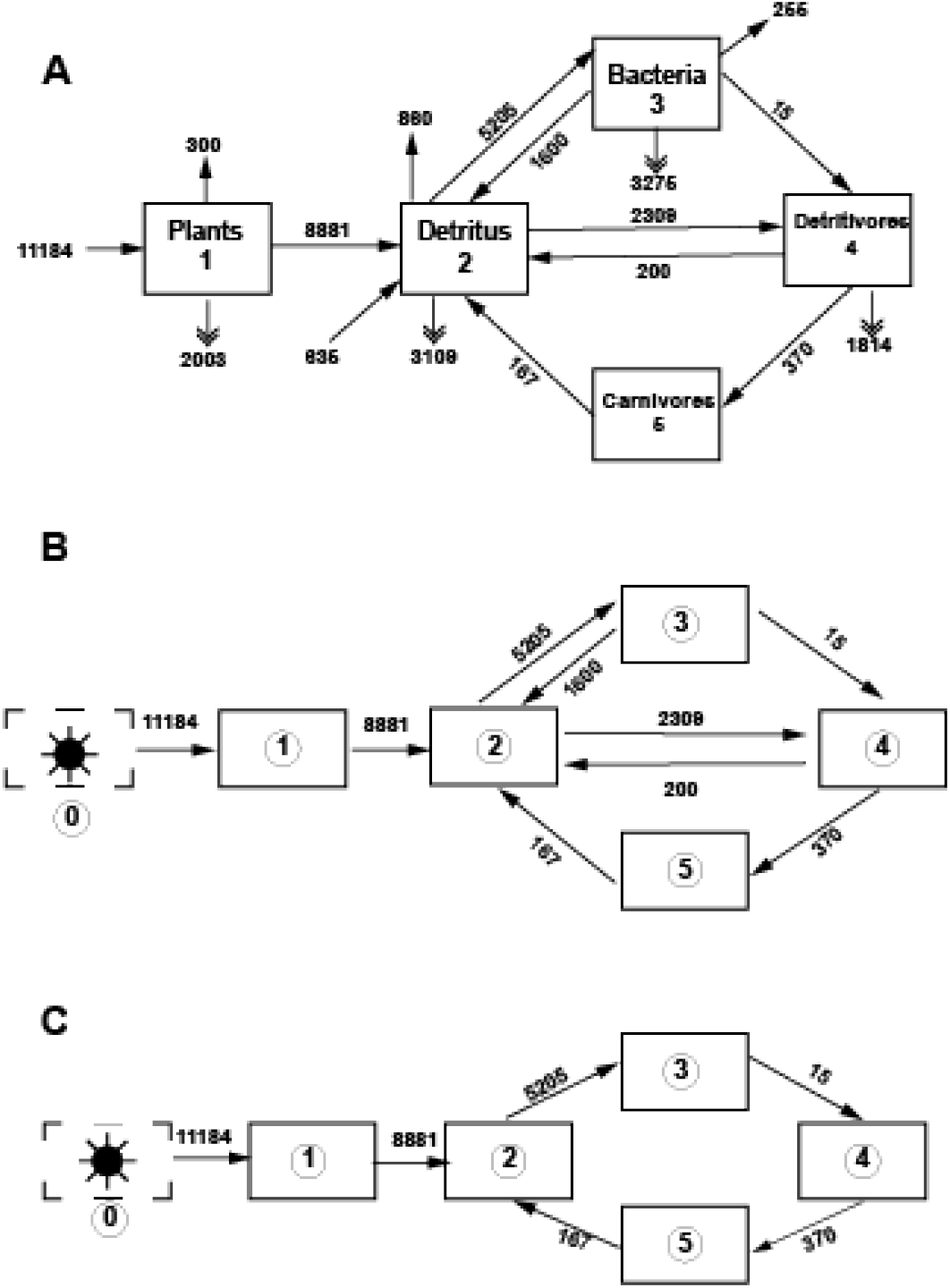
A) Example of the Cone Spring closed ecosystem as proposed by Ulanowicz (25). B) Simplified version of A considered in the estimation of the IV(t). C) Simplified version of B but in this case the flows between taxa 5 and 3 have been removed as well as the flows between taxa 2 and 5, and 3 and 2. The energy for taxon was estimated from the standing crop proposed by Tilly (40). Data and calculations are in Table S1.

The free energy principle states that any self-organizing system is at a non-equilibrium steady state with its environment and must optimize its (variational) free energy. We intend to define the notion of the Inner Value of an ecological network in these terms as a normalized amount of “free energy.”

We do know that there is extensive a controversial literature going back to the 80’s regarding the use of thermodynamic approaches to analyze the interactions within an ecosystem (i.e., 30-35). However, when considering an ecosystem, the only interactions that exist are via the transfer of mass or energy. That is the reality of the system and, having the possibility to estimate what is the degree energy/mass of exchange, there is no better way to calculate the conditions and the evolution of an ecosystem. Therefore, even though there are valid concerns, we consider that they are not strong enough to preclude the use of concepts like free energy or entropy (which are within the realm of the thermodynamic processes) to evaluate these interactions.

We use the notation drawn from (25, 41). In a system with n taxa, we have:

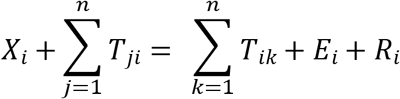

where X is the rate of solar radiation flow (for simplicity, we assume the existence of a single outside source of energy that affects a single taxon in the ecosystem and comes from a fictitious “taxon” 0); E_i_ is the rate of loss to the medium of taxon i to the outside world; Ri is the rate of dissipation of taxon i; T_ij_ is the rate of transfer from taxon i to taxon j, E_i_ and R_i_, for i = 1,...,n are left unspecified at this step. Then, the free energy principle could be expressed as (14)

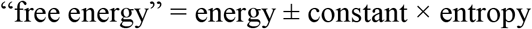

where energy is the expected value of some quantity of interest; entropy refers to a quantity measuring disorder, uncertainty, or complexity, and the constant translates between units of entropy and energy.

We interpret these magnitudes in the context of an ecological network N as follows:

- Energy (Total System Throughput): 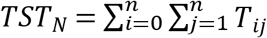
- Constant: the external input: *T*_0_=Σ_i_*T*_0*i*_
- Entropy: 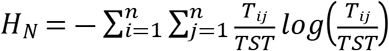

Then, the free energy is

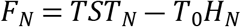

Notice that this expression is closely related to the notion of exergy, the goal function of ecosystems promoted by (37).

According to the free energy principle and, given the physical constraints on the taxa and their interactions, the actual FN of any given ecological network N at a steady state must be assumed optimal. Evolutionary forces use the free energy of the network to change its configuration by interacting with its environment. A network N is in a steady state when the system reaches a minimum value for F_N_.

By the definition of F_N_, we have that (since H_N_ ≥ 0) for any possible network N in the environment:

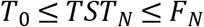

Notice that we assume that T_0_ is the same for all possible networks on the same class of taxa. Then, we have that 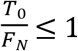. This allows us to propose a relative measure of the free energy of N as the Inner Value of the network:

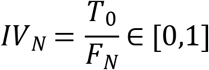

The rationale for this IV definition is that a system tends toward a configuration in which free energy is minimal and, thus, IV is maximal. However, this is a non-equilibrium state in which sudden changes in the environment may make it unstable, forcing a reconfiguration, at which the free energy is no longer minimal. One possibility is the unraveling of the system. This measure, unlike free energy, is invariant to constant changes in the values of the flows T_ij_, i,j = 0,...,n.

Considering a simple closed ecosystem (Fig. 1A) like the one proposed by Ulanowicz (25) for Cone Spring, we estimate, employing the scheme in Figure 1b, TST_N_, H_N_, and F_N_, resulting in IV_N_ = 0.15 (Table 1). The Inner Value can change if some flows disappear, yielding a network as defined in Figure 1c. In this case all variables are now giving IVN = 0.18 (Table 1). This indicates that the disappearance of some flows may decrease the inner value of the ecosystem. Another example is Chesapeake Bay network (38) flow model (Fig. 2) from which we estimated the same variables giving IV_N_ = 0.12 (Table 1). All data for these calculations are in the Supplemental Material (Tables S1 and S2).

**Figure 2.**
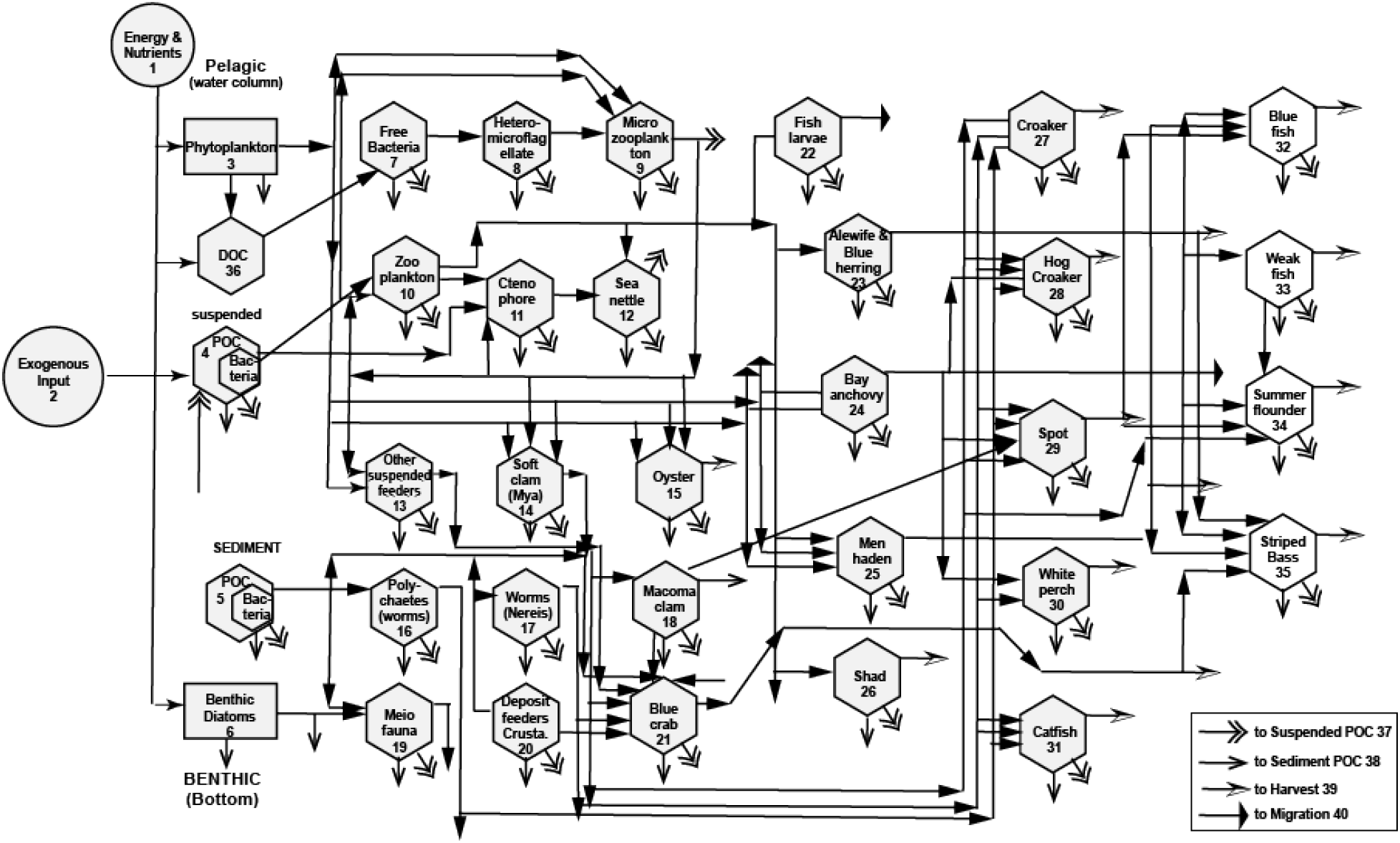
Diagram of the carbon flows of mid-Chesapeake Bay ecosystem modified from Ulanowicz (42). All values are in the Table S2

**Table 1.**
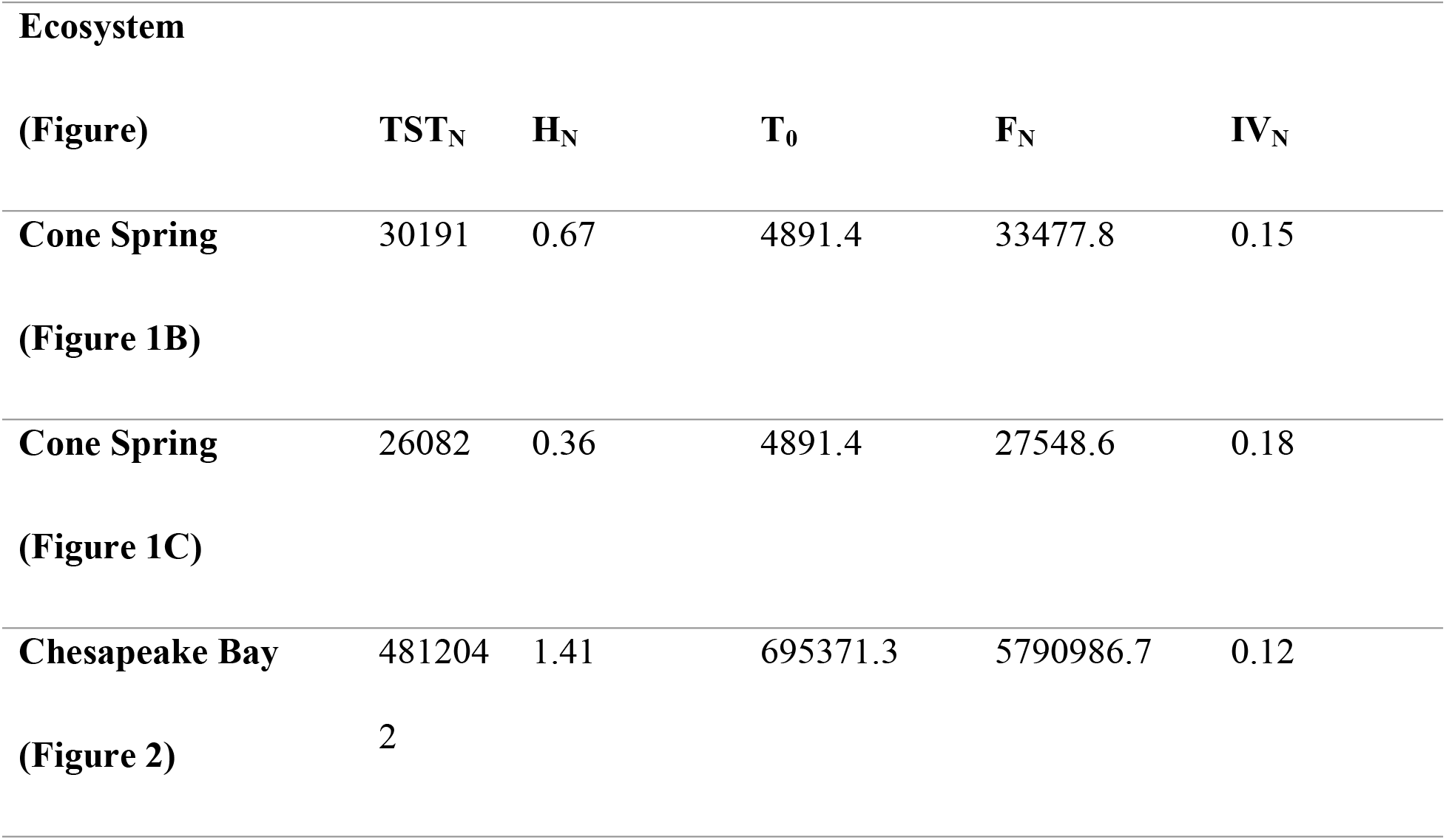
Estimation of the Inner Value for the examples in Figures 1 and 2.

Since we seek to assess the inner value of the system based on F_N_, we consider that IV can be an adequate variable to assess the time evolution of the ecosystem. Nevertheless, we do not provide, at this stage, a criterion about what is the actual health of the ecosystem based on the IV. That will require the development of a model to be test against ecosystems in different health stages.

However, we do propose an ideal evolution (Figure 3) in which the network starts to grow up from an initial configuration. It increases its size and complexity until it reaches a final period t_e_ of expansion or a particular threshold is crossed that further growth is not possible. It achieves then a metastable state while still being subject to both seasonal (i.e., regular) and random fluctuations. At that stage, it remains at a plateau, until at some time td the internal interactions make it susceptible to destabilizing forces that lead to its decay and eventual disappearance. Its inner value changes accordingly. This behavior is usual in open systems, whose internal states minimize free energy (41). Although for this ideal example we considered a continuous evolution, we assume that eliminating or adding taxa may produce jumps on the curve.

**Figure 3.**
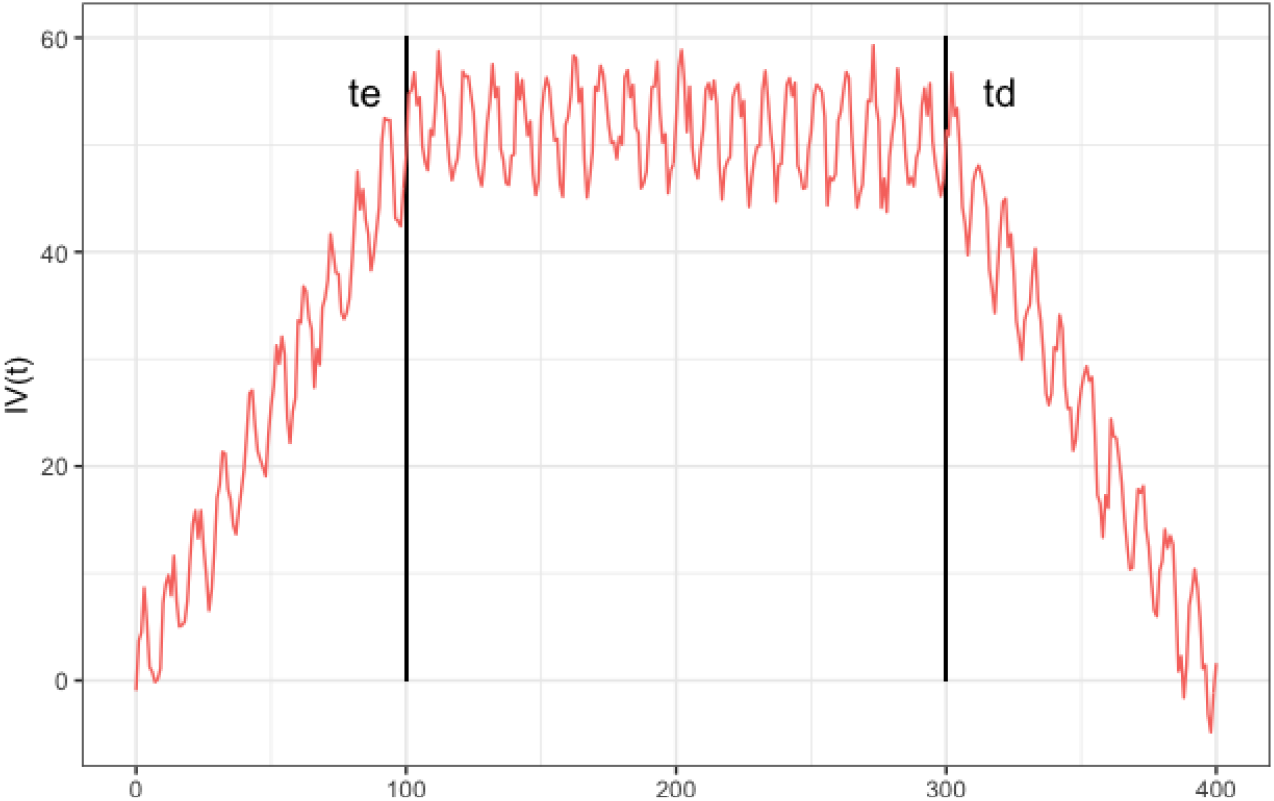
Schematic, time evolution of a closed ecosystem having three stages: grow, meta stable and declining until disappearance, with both periodic and random variabilities. Each stage is constrained by a threshold value (t_e_ *t_d_*). Although this is an ideal complete evolution, an ecosystem may have different behaviors (i.e., never reaching *td; continuous growth, etc*.). The IV is well suited to monitor such evolution

The notion of the Inner Value of an ecological network presented here is not the only measure that can be defined purely in terms of the internal flows of the network. For instance, Ulanowicz (25) summarizes a line of research into this topic based on Information Theory, presenting the concept of Ascendency.

Ascendency gauges the performance of the ecological system in processing its inputs. Its formal expression is

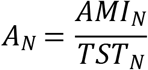

where AMI_N_ is the average mutual information of the system, relating the sources and the targets of flows, it measures how orderly and coherently the network flows are connected. The similarity between A_N_ and IV_N_ resides in the fact that both the former and F_N_ (the free energy of the network) yield a proportion of TST_N_. Ecosystems tend to increase both magnitudes (25).

However, perhaps more relevant are the differences between A_N_ and IV_N_. The first one is that AMI_N_ is just the contribution to H_N_ arising from local interactions, disregarding the chain connections among different taxa along the network. On the other hand, while A_N_ yields an absolute measure, IV_N_ is defined in relative terms, facilitating the comparison among different systems.

Exergy (37) is an alternative that can be defined as

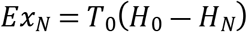

where H_0_ measures the entropy of the ecosystem’s environment; thus, Ex_N_ gauges how far the ecosystem is from achieving equilibrium. Then, in comparison with IV_N_, it has some disadvantages. The first one is defining the organization of factors external to the ecosystem. Another drawback is that different networks can be compared only in terms of how far they are from their equilibrium states but not their capacity to do practical work.

## Discussion

Ecosystems have an efficient organizational structure in which every member (taxon) has a clear, definite role. To maintain an efficient structure, every taxon has an adequate amount of free energy balanced throughout the ecosystem. Periodic changes (i.e., blooms) in a specific taxon rapidly trigger the response of dependent taxa in the trophic chain to keep the internal balance.

We introduce a novel approach over those proposed originally by applying the free energy concept to estimate the inner value (22, 25). In this sense, we cover the whole energy spectra by considering both the total energy and its entropy. By maximizing the latter, we ensure that the ecosystem is in its most stable condition, whereas the total energy provides an integral estimation of the system’s state.

When we analyze the ecosystem rationale, the system does not look for profit but for survival and expansion. It tends to develop services for external actors to help them achieve their goals. Also, ecosystems tend to be inherently and internally efficient and thus achieve the maximum output towards sustainability and growth. Inefficient branches in ecosystems are eliminated, and the rest of the ecosystem absorbs their mass/energy. Inefficient branches receive more energy than the one they return, generating an unbalance in the energy exchange, accumulating more internal energy in the node. The lack of frequent inner evaluation of the whole system may preserve these inoperative branches that consume energy that necessarily is extracted from nodes that generate an energy surplus. Therefore, the estimation of its inner value allows for defining the status of the ecosystem. On the other hand, if the Inner Value could be normalized and assessed across many different ecosystems, it may be possible to identify the status of the ecosystem. By analyzing its evolution through time, this value may give indications of the health of the system and what is its potential for growth or survival. Furthermore, the IV evolution could also indicate how the ecosystem behaves under some external stress.

Ecosystem services are a human-centered concept in which humanity receives some product (either cultural or commercial) or indirectly benefits from the ecosystem’s regulatory functions. In pursuit of extracting the most benefits from the ecosystems, humans have directly exploited almost all ecosystems on our planet or even modified them to obtain increasing profits. As we propose measuring the inner value of an ecosystem, we pretend to contribute to elucidating the key question Armsworth et al. (4) introduced: will we achieve greater conservation success by protecting nature for its own sake or our own sake?

In summary, estimating the Inner Value of an ecosystem could be a necessary requirement for any Environmental Impact Assessment (EIA). By modeling the changes in the inner value by the impact of any disturbance on a specific taxon or even the whole ecosystem, one can appreciate the degree of resilience the ecosystem may have, and one could define the maximum level of disturbance the ecosystem can support.

Modeling the inner value is also a factor when analyzing the service the ecosystem can provide and how much it can be extracted from it without crossing a threshold that may affect irreparably the ecosystem itself but also the service proper. As is the case of many polluted or disappeared ecosystems, the previous modeling could have prevented exceeding the damage and saving the economic loss due to the lack of the service(s).

In the present study, we propose estimating the inner value of an ecosystem as a tool to be applied at different stages and specific conditions to evaluate its health. Before any exploitation or anthropic disturbance that is bound to affect the ecosystem, estimating the inner value at the initial stage provides the basal level upon which to compare the impact of the disturbances may have in the future. However, also, it can be employed to model how different impacts, individually or acting simultaneously or sequentially, may have on the ecosystem. While we are discussing natural ecosystems, any economic system can also be regarded as an ecosystem where the agents interact among them, but also some of them must have relationships with external influencers (i.e., buyers, providers, competition). The measure of Inner Value can also be applied to establish the level of efficiency of an economic system when it is disconnected from external influences.

